# Black women undergraduates’ perspectives on how mentors and role models shape their beliefs about the attainability of science research careers

**DOI:** 10.64898/2026.06.15.731690

**Authors:** Wischell Joseph, Erin L. Dolan, Trevor T. Tuma

**Author notes:** **Author for Correspondence:** Trevor T. Tuma, University of Georgia, Department of Biochemistry & Molecular Biology, B202 Davison Life Sciences Building, Athens, GA, 30602.

## Abstract

Undergraduate research experiences offer important paths into scientific research careers, yet students do not experience them uniformly. For Black women, these experiences occur within racialized and gendered environments that may shape whether they perceive research careers as attainable. Yet little is known about these influences, including how mentors, role models, and institutional contexts, could support or limit Black women’s beliefs about research career attainability. To advance our understanding of these influences, we conducted interviews with 23 Black women who participated in undergraduate research at 18 institutions in the United States, including historically Black colleges and universities (HBCUs) and predominantly White institutions (PWIs). We conducted qualitative content analysis to understand Black women undergraduate researchers’ perceptions of the attainability of a scientific research career, including the influences of their mentors, role models, and institutional context. Three main themes emerged. First, Black women undergraduates varied in the importance they placed on sharing racial and gender identities with mentors and role models; some viewed such similarities as highly meaningful while others described them as less influential. Second, Black women undergraduates described that mentors and role models who shared similar life experiences, values, attitudes, or beliefs contributed to perceptions that scientific research careers were attainable, regardless of gender or racial similarity. Third, institutional context (HBCU, PWI) shaped how mentoring and role modeling influenced Black women undergraduates’ perceptions of research career attainability. We conclude by offering recommendations for individuals seeking to support Black women in undergraduate research and in their pursuit of research careers.

## INTRODUCTION

The continued economic prosperity and scientific advancement of the United States will depend on its ability to develop a qualified and diverse scientific workforce ^1^. Although women have made gains in participating in biology, racial and gender disparities remain prevalent within many STEM fields. In particular, Black women remain underrepresented in many science-related research careers despite longstanding efforts to broaden participation. These disparities do not reflect differences in talent or motivation, but instead reflect structural inequities that shape minoritized students’ experiences in science ^2,3^. Across K-12 and higher education, Black students contend with systemic barriers including unequal access to resources ^4,5^, stereotype threat ^6^, and microaggressions ^7^.

For Black women in particular, these experiences have the potential to be influenced by both racism and sexism in ways that are distinct from those of White women, Black men, and other minoritized groups in science ^8^. This experience has been characterized as a “double bind,” in which Black women must navigate both gender and racial biases that compound one another and create unique barriers that limit their ability to thrive ^9^. These barriers can negatively affect Black women’s persistence in STEM degrees and discourage their pursuit of scientific research careers ^5,10^. Even when Black women remain in science, they continue to encounter ongoing marginalization, such as tokenism ^11^, isolation ^8^, disproportionate service expectations ^12^, and the “pet-to-threat” phenomenon (i.e., initial praise and visibility followed by perceptions of threat once their competence or influence is realized) ^13^. Understanding how Black women come to view research careers as attainable is critical for broadening participation in the scientific workforce.

Undergraduate research experiences function as important steppingstones in the path towards a career in scientific research. Participation in undergraduate research supports the development of technical skills ^14,15^, cultivation of a scientific identity ^16–18^, formation of professional relationships ^19^, and sustained or increased interest in graduate education in science ^14,15,20^. Within these environments, more experienced scientists (e.g., faculty, postdoctoral research associates, and graduate students) often serve as mentors and role models who support undergraduates’ integration into the scientific community ^21–23^. Undergraduates who receive greater mentorship during undergraduate research report gains in their scientific self-efficacy, science identity, motivation, and scholarly productivity ^19,21,24–26^. However, access to research experiences, mentorship, and role models alone does not guarantee that students will view research careers as desirable or attainable. Students’ experiences within undergraduate research can vary not only by program structure, but also by the quality of their relationships with mentors and other influential individuals in their research labs ^27–29^. Although undergraduate research and mentorship are often position as key educational interventions for supporting persistence in science-related careers, the extent to which these experiences shape students’ career trajectories depends on how students interpret and experience them ^30^. In particular, interactions and support from mentors may shape whether students come to view a scientific career as accessible, realistic, and personally attainable.

Considerable scholarly attention has focused on understanding what characteristics make mentors and role models influential for students. One commonly studied factor is perceived similarity, or how similar a student feels to their mentors or role models. Many educational initiatives and diversity efforts have emphasized demographic matching (e.g., shared gender or race/ethnicity) under the assumption that students benefit most from mentors and role models who share their gender or racial identities. This assumption is grounded in theory suggesting that individuals are more likely to attend to, identify with, and emulate others whom they perceive as similar to themselves ^31–33^. Consistent with this, women and individuals from racially minoritized groups express greater preference for a mentor who shares similar demographic backgrounds ^34–36^. However, some research on the effects of demographic matching shows small effects ^35,37–39^. Other research shows mixed effects or no effects at all ^40–42^. Similarly, research on STEM role models shows variable effects of demographic matching on students’ motivation and persistence, with studies reporting positive, null, and sometimes contradictory findings ^43–47^. These results suggest that demographic similarity alone may not always be sufficient for explaining how these relationships benefit students and shape their experiences.

Students can perceive similarities with mentors and role models in other ways beyond demographic characteristics. For example, students may perceive sharing similar perspectives, values, and outlooks with their mentor (i.e., psychological similarity)^48^ or sharing similar education, career, or life experiences (i.e., experiential similarity) ^49^. Importantly, students who perceive greater psychological similarities with their mentor report more positive relationship outcomes than those who do not ^39,41,42,50^. These trends are robust across sociodemographic backgrounds, suggesting that psychological similarity is impactful for all students ^39,41,42^. These findings raise important questions about how Black women undergraduate researchers’ similarities with their research mentors and role models influence their perceptions that research careers are attainable.

These findings also underscore the need to better understand how contextual factors influence Black women undergraduate researchers’ experiences. Institutional environments can play a central role in structuring students’ access to undergraduate research, the availability of mentoring relationships, and the degree of representation they encounter in STEM. Institutions of higher education vary substantially in the gender and racial composition of both students and faculty, as well as in the extent to which students experience inclusion, support, and belonging ^51^. These differences may be particularly consequential for Black women as they simultaneously navigate racialized and gendered environments. One contextual factor that may shape these experiences is the institutional setting, particularly whether students are enrolled at predominantly White institutions (PWIs) or historically Black colleges and universities (HBCUs). The impact of similarity to a mentor or role model may vary depending on whether such forms of similarity are common, rare, or institutionally reinforced. For example, same gender or same race mentoring relationships may have different effects in environments where Black individuals and women are highly visible compared to contexts where they are more isolated. The institutional context of an HBCU or PWI may shape not only who is available to mentor, but also what those relationships signal about the attainability of a research career. Relatively little research has examined the role of the institution in shaping Black women’s undergraduates’ research experiences, research mentors and role models, and their subsequent influence on their beliefs about the attainability of a research career.

### Present study

We examined how Black women undergraduate researchers’ perceptions of their research mentors and role models and their institutional context (i.e., predominantly Black or predominantly White institution) influenced whether they viewed research careers as attainable. To accomplish this, we conducted a qualitative study to understand the experiences of Black women undergraduate researchers and how they perceive the attainability of research careers. We used an exploratory approach with inductive reasoning to identify patterns in the narratives and lived experiences of students, their perceptions of the ways in which their mentors or role models shaped their views of research career attainability, and their perceptions of how institutional context was influential, if at all. As a Black woman student researcher and former HBCU student doing research at a PWI, the first author brought an insider perspective to this work.

## METHODS

### Ethical guidelines

All participants were treated in accordance with APA ethical guidelines. This research was conducted with approval from the University of Georgia’s Institutional Review Board (PROJECT00010581).

### Participants and Recruitment

We recruited students who identified as Black women and had conducted undergraduate life science research for at least three months to participate in the study. To recruit participants, we directly emailed undergraduate research program directors, program leaders, and advisors from PWIs and HBCUs primarily in the southeastern and mid-Atlantic United States and asked them to forward our study invitation to eligible students. Participants were also encouraged to share the study information with peers and friends whom they believed met the study criteria. The study invitation contained a link to an online screening survey administered through Qualtrics. In the screening survey, participants self-reported their eligibility, provided contact information, and completed a brief demographic questionnaire. We reviewed screening survey responses on a rolling basis, and eligible students were invited to participate in interviews.

We strategically recruited students from both HBCUs (defined as institutions with less than 15% White students enrolled) and PWIs (defined as institutions with less than 15% Black students enrolled) to capture the experiences of students across differing institutional contexts. We also recruited strategically to ensure that the participant pool reflected variation in Black racial and ethnic backgrounds (e.g., African American, Afro-Caribbean, African, Afro-Latino, Black, Haitian), life science majors (e.g., biochemistry, ecology, neuroscience), and highest level of parental or guardian education. Participants also reported a variety of career aspirations, post-baccalaureate degree aspirations (e.g., pursuing MD, PhD, MD-PhD, and MPH programs), and were moderately to extremely interested in pursuing research careers. A total of 33 students met the study criteria and completed the screening survey. Of these students, ten declined to participate in an interview, resulting in a final sample of 23 students representing 18 institutions. Of these, students represented 12 distinct HBCUs and 6 PWIs. Additional demographic information for the participants who completed interviews is presented in Table 1.

**Table 1:**
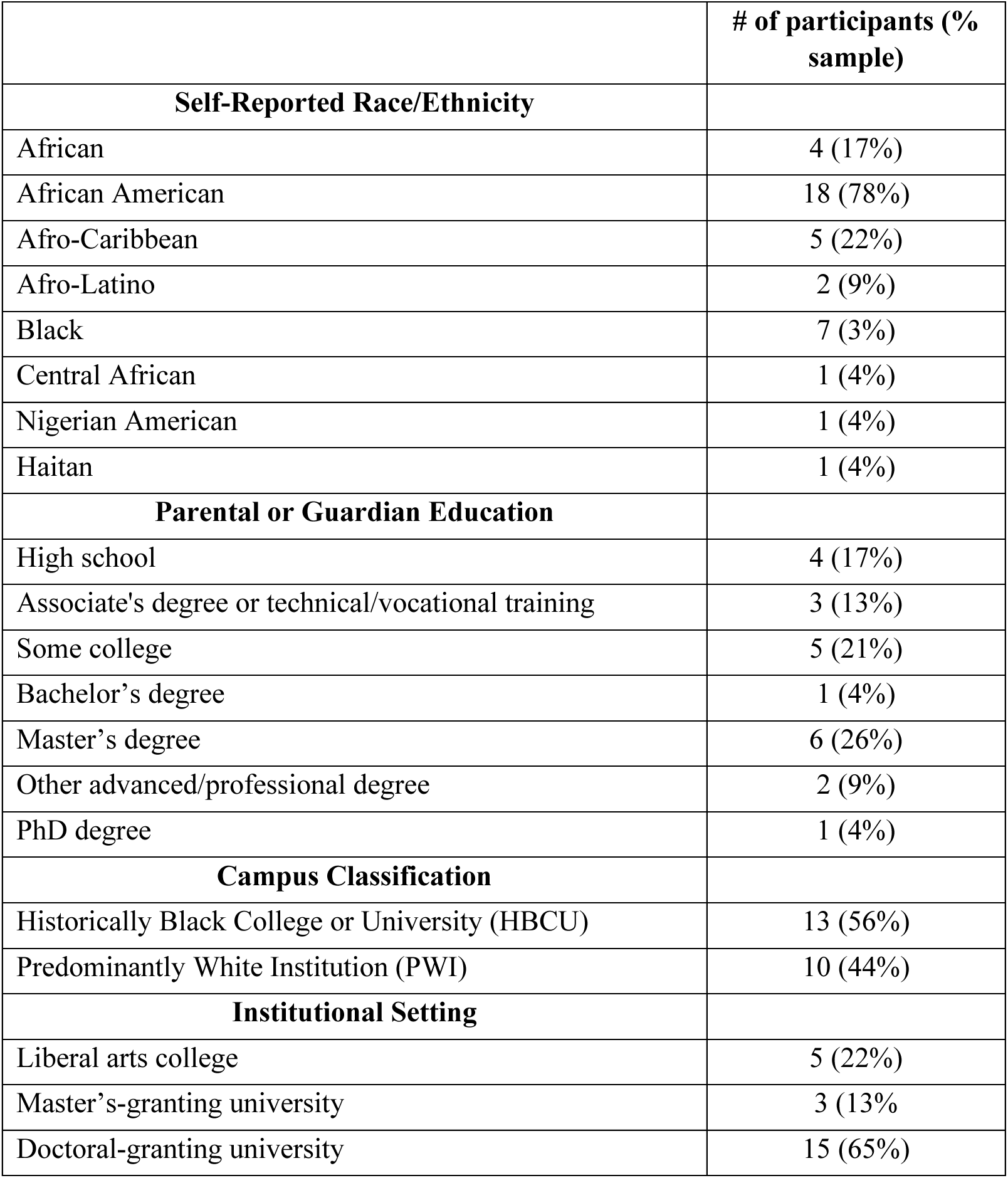
Participant demographic information (*n* = 23).

### Data Collection

The first author interviewed the participants on Zoom using a semi-structured approach. The interview protocol asked students to describe their mentoring experiences during their undergraduate research experiences, including who influenced their career decision making and the role of mentors and role models in this process. Participants were also asked to describe the influential individuals within their research labs, the characteristics and behaviors that made these individuals influential, and how these relationships shaped their views of their ability to be successful. Additional questions explored how their mentors’ race and gender influenced participants’ research experiences and beliefs about whether a research career was attainable, if at all. Participants were also asked to reflect on their broader experiences as Black women conducting undergraduate research at their institutions and how their experiences attending a PWI or HBCU shaped their career aspirations. Interviews lasted an average of 45 minutes (range = 30-60 minutes). Participants received a $30 Visa gift card as compensation for participating in an interview. All interviews were audio recorded and transcribed verbatim. We reviewed transcripts for accuracy and removed or replaced identifying information (e.g., names, institutions) with pseudonyms.

### Data Analysis

We analyzed the interview data using an iterative inductive-deductive approach to thematic analysis ^52^. We began by reading each transcript individually with our research question in mind and writing analytic comments about recurring ideas, patterns, and notable experiences described by participants. We engaged with relevant literature to consider how concepts from prior research aligned with emerging patterns. For example, participants’ responses reflected ideas related to identity salience, intersectionality, mentoring relationships, and belonging in STEM. We used this literature to inform our developing interpretations while also remaining grounded in participants’ recollections. We then used inductive open coding to examine how participants described their mentoring experiences, influential individuals in their research labs, and perceptions of how the race and gender of their mentors and role models shaped their research experiences and career aspirations. Because we intentionally sampled across institutional contexts (i.e., HBCUs and PWIs), we also attended to how participants described navigating different institutional environments and how these contexts influenced their experiences.

Throughout coding, members of the research team repeatedly read and discussed transcripts to refine our interpretations and compare patterns across interviews, clarifying relationships among codes ^53^. As codes evolved, we revisited transcripts and reviewed excerpts associated with each code to better define distinctions among codes and compare how similar ideas were expressed across participations. In a final stage of analysis, we grouped related codes together to develop broader themes that captured shared patterns in participants’ descriptions of their mentoring experiences, representation, institutional context, and the attainability of research careers. All themes were discussed amongst the researcher team and refined until consensus was reached. Rather than calculating interrater reliability, we discussed differences in our coding through discussion to ensure rigor and develop a shared understanding of participants’ experiences ^54^.

### Positionality

The research team consisted of three researchers who, at the time of the study, were a post-baccalaureate researcher, a postdoctoral research associate, and a faculty member working within a life science department. The researchers held differing social identities and professional experiences. The first author is a Black woman who had recently completed her bachelor’s degree and was contemplating a career in biomedical research. Her lived experiences navigating STEM and research environments informed the interpretation of participants’ decisions surrounding career decision-making in the life sciences. The other two authors are White, each holds a doctoral degree in biology, and are biology education researchers whose scholarly interests focus on career development and mentoring relationships in STEM research. They have work closely with and mentored students from a range of backgrounds, including Black and African American students, throughout their careers. During some of the interviews, the interviewer disclosed aspects of her identity and shared how her own experiences resonated with those shared by the participants. The study design, data collection, and interpretation of the findings were shaped through an ongoing collaboration between the coauthor’s lived experiences and the broader scholarly expertise of the research team.

## RESULTS

### Black women undergraduate researchers vary in the importance they place on racial and gender similarity with their mentors and role models

Students in our study highlighted how mentors and role models who shared their identities as Black and/or women shaped their beliefs that a career in science research was attainable. Some students noted the importance of sharing both identities with their research mentor or role model. For example, one student described how sharing both racial and gender identities with her mentor made the mentor’s accomplishments appear more attainable. She explained, *“She is a Black woman like myself. Seeing her make strides in the field inspires me, and it makes me more confident that I can do the same thing that she is doing.”* Other students lamented not having access to a Black woman who could be a mentor or role model. They felt that working with a Black woman principal investigator (PI) would be empowering and make it possible to see themselves in this role, as this student explained:

> *“Just because he’s a white man, I wouldn’t say I could see myself in his role because it’s like a different identity. I think it definitely would feel a bit empowering if my PI was a Black woman. I still think he’s like helpful and he’s provided a lot of guidance, but I couldn’t picture myself in his role because we don’t share like the same identity… I definitely don’t see too many other Black women researchers… I think the absence of Black women in research roles has affected my perception of pursuing research as a career path.”*

Although this student acknowledged that her PI (i.e., a white man) provided helpful guidance (i.e., mentoring), the absence of shared identity limited her ability to imagine herself pursuing a research career (i.e., role modeling). This disconnect was mentioned repeatedly, with another student noting, “*I think if any of the people I identified had looked like me, I would’ve been more comfortable exploring other career paths in research.”*

One student noted that the collective identities of her role models made a research career seem attainable. This student knew a successful woman researcher who was not Black and a successful Black man researcher, and she considered both together in viewing a research career as attainable, as she explained,

> *“Since my PI at [University] was female it was like ‘OK, we got this,’ and since my research advisor here is Black but he’s male so it’s like, ‘OK I’m just a mixture of both, I can do this.’”*

Some students described that having a mentor or role model who was Black helped them feel more comfortable, welcomed, and seen, regardless of the mentor’s gender identity. One student explained that her mentor’s race “makes a difference” but her gender did not because it was easier to find women in the field. Another student who had a Black male mentor described how his being male didn’t make her feel less capable of being successful in a research career because “*there’s a lot of females working there. So, I never felt like he had a better position because he was a male.”* However, the fact that he was Black was important because she had not seen *“a lot of Black people in the lab.”* She noted that observing him and his success helped her see how she could succeed. She described, *“being able to see him doing so well, and him specifically being my mentor I felt like I could see myself in him.”*

Other students emphasized the importance of their role models being women, regardless of their racial identity. One student explained how her research mentors were men and she did not see them as role models because of this, but other professors were role models because they were women:

> *“Dr. [Professor] inspires me since she’s a woman. She’s one of the most respected professors at my school. The way she teaches, does research, people know her and respect her. For my PI and bench mentor, not so much, because they are men. Dr. [Professor] (has) been an incredible role model to me. Seeing her thrive and succeed has shown me that, even though she’s white, she’s a woman and she’s thriving.”*

Other students emphasized that having a woman mentor helped them develop a comfortable working relationship. For instance, one student described feeling *“more comfortable talking about personal stuff with her [woman mentor] and building more of a personal relationship as compared to my male research mentor.”*

Not all students perceived that sharing gender and racial identities with their mentors or role models was necessary to pursue a research career. Some found that doing research in a diverse lab environment with high-quality mentorship made them feel that a research career was attainable regardless of the race or gender of their mentors or role models. One participant with an Asian mentor stated, “*I didn’t feel like an outsider… I still got treated with the same respect as anybody else.*” Some students noted that the diversity of their research group mitigating any feelings of “outsider” status.

Another student emphasized the importance of positive relationships for engendering a feeling of belonging and a positive research experience. She explained, “*you can have a great personal, professional connection with people that don’t look like you, and still have a great experience and hone in on what you need and gain what you need as a person of color.”* These examples show that, although shared identity was powerful for some students, others felt that being treated with respect and experiencing supportive relationships was what they needed to see a research career as possible.

Racial and gender similarity did not compensate for the negative effects of poor mentorship. One student described how they were initially excited to work with a Black woman mentor for the first time, expecting to find an ally but how that did not turn out to be the case, *“She kind of gave the impression that she wants everyone to know she’s the smartest person in the room… she was just really mean… I was like ‘I’m not gonna get a good letter’ and she was just like ‘you’ve been doing an awful job’.”* This experience was unexpected and left the student feeling unsupported and unwelcome, undermining her confidence at a critical point in her pursuit of a research career.

Finally, several students in our study explained that they believed a research career was attainable primarily because of their own attributes or effort rather than because of any mentor or role model. They described how gaining experience, putting in time and effort, and being interested and confident were important for a research career to be attainable, as this student explains:

> *“I feel as if I can be successful because it just really goes on your skills. If you can do the work and do it effectively… You have to be confident too. That’s why I learned a lot, too. You have to be confident in what you’re doing and you should be fine…”*

### Black women undergraduate researchers shared experiences and values with their mentors that made research careers seem attainable

Some students who described racial and gender similarity with their mentors and role models as unimportant highlighted how shared experiences were influential. Students framed their experiential similarity, meaning their shared lived experiences, in terms of being an “outsider,” such as being marginalized or minoritized in some way or otherwise encountering and overcoming hardships. One student explained that sharing a non-dominant background with her mentor convinced her a research career was attainable, *“It wasn’t more someone who looked like me but more like someone who comes from a similar background as me, like an immigrant background. It made me realize you don’t have to be a white American man to do research and be successfu*l.” Another explained how commonalities in their background made their journey more relatable and thus attainable. For instance, one student described how her mentor, *“told me she was once in my shoes… she faced many of the same hardships I did and still came out very successful. That made me feel like I could do it too.”*

Other students highlighted how sharing values (i.e., psychological similarity) with their mentors and role models was influential. Students observed how their mentors and role models embodied their values of achieving some balance between work and life and of making a difference in the world. Several students had considered careers in medicine but saw them as demanding and inflexible. These students saw a research career as a healthier and more sustainable lifestyle. One student recalled a candid conversation with her mentor, who was a staff scientist: “*He has his PhD. He just works there… His work-life balance is beautiful. My mentor is the reason why I picked PhD, for sure.”* Others echoed this sentiment, citing research as an impactful yet balanced career. One student stated, “*I want a good work-life balance, and research offers that*,” and another student note that research “*is impactful in a different way*.”

One student noted that the absence of shared values with their mentor was a deterrent to pursuing a research career. Seeing their mentor and other researchers overwork made a research career unappealing. She explained about her mentor, “*With him (there) not a lot of work-life balance. There were days where he’d forget to eat… One time we were in lab until 9:00 p.m., and he didn’t even realize his glove was burning, and we were working over a Bunsen burner. He said it took him like 30 seconds to realize.”* The same student also recalled a lab member keeping a toothbrush and mug in the lab, as if they lived in the lab. She remarked, “*I’m like, ‘You’re brushing your teeth here? What are we doing?!’*”

### The environment influenced the salience of racial identity for assessing attainability of research careers

For the Black women in our study doing research at a PWI, being one of few or the only Black woman in their research lab heightened awareness of their identity and dampened their perception that a research career was attainable. One student explained how the absence of “*people who looked like her*” made it difficult to see herself in a research career:

> *“If you don’t see yourself represented, you almost feel like you can’t do it because you don’t see people in that position. Even the one (Black woman) biology professor I mentioned, she’s not a researcher, she’s just a [lecturer]. So there are no Black female researchers, specifically in the biology department at PWI.”*

Another student echoed this sentiment, emphasizing not only that it was important to see people who looked like her as role models, but also that talking with Black women researchers would allow her to have more informative conversations about this career path:

> *“I think if any of the people I identified had looked like me, I would’ve been more comfortable exploring other career paths in research, because I’d literally be able to see myself doing that. I’d have been more open to researching it, looking into it, thinking about it more. I would’ve had someone I felt comfortable talking to about my concerns like, ‘This is what I’ve always dreamed of [referring to a career in medicine], but now I’m seeing this other path [referring to a career in research]. Can we talk about it?’”*

One student emphasized the racial and ethnic diversity of her lab environment rather than her institutional environment for influencing her views about research career attainability. She noted that being a part of a diverse lab group made being the only Black person less remarkable. She elaborated that she was *“the only black person that was accepted to the internship, but I didn’t feel that in the lab because it was a pretty diverse lab. My lab partner was from Jordan. Another lab member was from Brazil so it was a pretty diverse community. I didn’t feel out of place.”*

Some students perceived their institutional context as important for gaining experience they viewed as critical for attaining a research career. Students who attended HBCUs and did research at PWIs in the summer highlighted that the size and research intensity of the PWI offered more varied opportunities to do research. One student explained about her research experience at a PWI, *“there was more in terms of like having more opportunities to research, it doesn’t even have to be professors but like having more opportunities to say, ‘Oh, I want to research in this field,’ and I can actually do it.”* She elaborated that the smaller size of her HBCU, which was home to fewer faculty, did not afford the same breadth of options:

> *“There’s specific research that some specific professors do. If you don’t align with that, or if you can’t really work with them in the lab, then you can’t really do research at all. I wish that was different. I also wish that my school kind of encouraged us to pursue more like external research, like summer research, like apply for more research opportunities outside of the state.”*

Some students at HBCUs felt at a disadvantage in attaining a research career because graduate programs were looking for more extensive experience doing research at the undergraduate level, which they felt was not possible at their institution. This student explained:

> *“Since we’re smaller, we don’t have a lot of research experiences. I have been blessed to be able to do research at my institution but I wasn’t able to start that until my junior year. I know a lot of programs now look for people to have published and those things take time… We don’t have a lot people coming in as freshmen doing research. Having the opportunity to do (research) all the way through undergrad - that’s unheard of.”*

Considered together, the students in our study attended to aspects of their institutional environments that afforded or constrained their perceptions that research careers were attainable. Some focused on the racial makeup of their institutions or research groups, while others were more mindful of how their environments influenced the type and amount of research they could do.

### Limitations

We encourage readers to consider several limitations of our study. First, our study focused solely on Black women who had engaged in life science undergraduate research and who expressed interest in research careers. This design missed the experience of students who were unable to secure a research experience or became disinterested in research. Second, our results relied on data collected at a single timepoint from a relatively small number of Black women. This design afforded richer insights for generating hypotheses about how Black women’s mentors and role models influence their views of research career attainability. Longitudinal research following larger samples of Black women as they conduct undergraduate research and make career decisions would help clarify students’ evolving perceptions of research career attainability and how mentors and role models influence this process. Finally, our study centered the narratives of Black women in life science fields. This is a strength because it afforded an in-depth understanding of Black women’s views of research career attainment in the life sciences. Yet our findings are not necessarily transferable to the experiences of Black women in other science disciplines (e.g., chemistry, physics, engineering) or to individuals with different intersecting gender and racial identities (e.g., Black men, Black non-binary students, Hispanic women).

## DISCUSSION

In this study, we sought to explore the characteristics of mentors and role models that Black women undergraduate researchers perceive as influential for viewing research careers as attainable. The students in our study held mixed views of the importance of racial and gender similarity with their research mentors and role models. Mixed views may explain the absence of any clear pattern in prior research regarding the importance of demographic matching. Our findings suggest several interrelated phenomena may be at play.

First, students who spoke about the importance of racial and gender similarity emphasized how individuals who shared these identities helped them perceive research careers as attainable. Regardless of whether these individuals were their research mentors, the Black women in our study spoke about their influence as role models (i.e., mentors could be role models and role models didn’t have to be mentors). Their perspectives indicate that mentoring and role modeling are separable forms of support, and that racial and gender matching may be more important for role modeling than for mentoring. This interpretation is consistent with the notion that observing role models and assessing whether their position seems feasible or attractive are processes through which individuals take on new professional identities ^55^. It is also consistent with prior research showing that role models can function as inspiration, providing exposure to possible careers without the direct support characteristic of mentoring relationships ^26^. This is encouraging, as it suggests that such influence could be achieved at scale without the time and energy required to engage in a meaningful mentoring relationship. Future research should aim to further delineate the influence of mentoring versus role modeling in research career development and decision-making, especially among students with racial and gender identities not well represented in these career paths.

Second, identity theory suggests that different identities are more or less active across situations ^56^. Identity salience refers to the likelihood that a given identity will be invoked across situations ^57^. The Black women in our study may have varied in the salience of their intersectional identity as Black women, as well as in the salience of their identities as Black individuals or as women separately. The situations of being in an HBCU or PWI or in a life science research environment likely functioned as cues (or not) that activated students’ racial identity. Indeed, some students in our study noted that gender identity was less important because of the prevalence of women in the life sciences. Other students noted their racial identity was less important because of the racial diversity of their lab group. Our findings suggest that identity salience, lab group diversity, and institutional diversity may function as moderators of the effects of shared race and gender identity in mentoring relationships and may help account for the mixed effects reported in the literature ^38,39^.

Students in our study also indicated that other forms of similarity with their mentors – namely sharing experiences and values – were important for perceiving research careers as attainable. This finding echoes existing research showing that individuals who perceive greater psychological similarity, such as shared values, interest, and beliefs, with others report higher quality and more effective working relationships ^39,40,42,48,58,59^. This has important implications for practice, because matching mentees and mentors on the basis of shared gender and/or race may not be feasible. However, there is evidence that interventions can successfully foster perceptions of psychological similarity ^60,61^, offering an alternative way to promote the quality of Black women’s mentoring relationships with mentors who may be demographically dissimilar from them. Mentees are more likely to develop a sense of psychological similarity with their mentors when the pair has opportunities to get to know one another and spend time identifying shared experiences, interests, and values. For instance, students in our study described how their mentors disclosed their experiences of being an “outsider” and this disclosure created a feeling that they had outsider status in common. It may be that mentoring professional development is effective at least in part because mentors learn tactics for opening up lines of communication with their mentees ^62^, thus creating space for a sense of similarity to emerge. In practice, mentors should be encouraged to use communications strategies to create space for mentees to help identify even small points of a shared perspective or common ground, as doing so has the potential to foster a productive mentoring relationship with Black women mentees.

Our results highlight that some Black women may prefer to have a mentor who shares their gender and racial background. If there are Black women with the interest and capacity to serve as mentors, then such preferences could be accommodated. However, our findings also highlight that such matching is not a prerequisite for a high-quality mentoring relationship or for perceiving research careers as attainable. Instead, students highlighted the potential of developmental networks ^63,64^ – in other words, having connections to a group of people who can provide different types of support and serve as role models for different identities. Some students described how they didn’t need to have a single person who was both Black and a woman to view a research career as attainable. Rather they described how their networks included women who demonstrated that success was possible and Black people (men) who demonstrated success was possible. This finding echoes prior research on the importance of developmental networks for STEM identity formation among undergraduate students ^65^. This also has implications for practice; students could be encouraged to reflect on their career interests and aspirations and consider a collection of people who could provide different forms of mentorship and role modeling ^66^. Mentors can also be reminded that they do not have to be a singular source of support for their mentees. This messaging is especially critical for Black women mentors given that women ^67–69^ and people of color ^70^ can bear a disproportionate share of academic service work, including formal and informal mentorship. In sum, the perspectives of the Black women undergraduate researchers in our study indicate that there are a variety of ways that Black women can be supported by mentors and role models in viewing research careers as attainable.

## Supporting information

Supplemental Materials_Interview Protocol

## ACKNOWLEDGEMENTS

We thank colleagues who assisted with distributing our study invitation, and we thank our participants for sharing their time and perspectives. This work was supported in part by funding from the Georgia Athletic Association Professorship for Innovative Science Education, National Institute of General Medical Sciences of the National Institutes of Health Award Number R25-GM109435, and National Science Foundation Division of Graduate Education Grant Number 2328692. The content is solely the responsibility of the authors and does not necessarily represent the official views of the National Institutes of Health or the National Science Foundation.

