## Supplemental Materials_Interview Protocol for "Black women undergraduates’ perspectives on how mentors and role models shape their beliefs about the attainability of science research careers"

1. Can you tell me about a typical day in your lab?
    a) How did you pick this research to work on?
    b) Do you remember why you decided to do research in the first place?

2. Take a minute and think of the person in your research group who is most influential in your research experience. This can be someone who provides you with guidance or is influential in other ways. This person can be a positive or a negative influence. Feel free to take your time and let me know when you have someone in mind.

3. Ok, do you have someone in mind?
    a) Tell me about this person. What is their role? How do you interact with them?.
    b) Why did you select this person as the one who is most influential for you?
    c) What are things they do that make them influential?
    d) How often did you talk with them? And what types of things did you talk about?
    e) How well do you feel like you know them? How well do you feel like they know you?

4. I am going to share 2 definitions, one of a role model and one of a mentor.
    A role model is defined as someone whose success demonstrates that achievement is possible. A role model provides an example of the attitudes and behaviors needed to achieve similar results. Role models can be someone you look up to but do not know or someone who you know well.
    A mentor is someone with more experience who directly helps and supports a less experienced individual, to enhance their personal and professional development.

With these definitions in mind, would you identify the person you're thinking of as a role model, mentor, or both?

Is there anyone else who you may consider a career role model?

If the participant says role model, ask this:
-What makes them a role model for you?

If the participant says mentor, ask this:
-What do they do to mentor you?
-What makes you think of them as a mentor?

5. Do you think of this person as successful?
    -If yes, do you think you can be successful like them?
    -If not, why not be successful?

6. Does this person inspire you to follow in their footsteps?
    - If yes, what about them is inspiring?
    - Probe about example(s) of traits this person has or things this person does that are inspiring, and why these traits and behaviors are inspiring.

7. Does this person's gender and race make a difference for you? Tell me about that.

Now I'd like to ask you about your career goals and whether this person has been influential. First, I'd like to learn more about your career goals. You have been working on research for some time now.

8. Do you think a research career could be the right path for you?
    a) Why or why not?

9. Has the person you have told me about influenced your thinking about whether a research career is right for you? Tell me about that.

10. Have your career goals changed since working with this person? Tell me about that...

11. What has your experience been like as a Black woman doing research at this school?

12. Have you ever talked about being a Black woman in research with them?

13. Have they done anything to show you that you're being valued?

14. Does this person's race and gender make you feel like someone like you could succeed in their role at this school?
    a) Why or why not?
